# Bacterial processing of glucose modulates *C. elegans* lifespan and healthspan

**DOI:** 10.1101/2020.09.13.295451

**Authors:** Samuel F. Kingsley, Yonghak Seo, Calista Allen, Krishna S. Ghanta, Steven Finkel, Heidi A. Tissenbaum

## Abstract

Intestinal microbiota play an essential role in the health of a host organism. Here, we define how commensal *Escherichia coli (E. coli)* alters its host after long term exposure to glucose using a *C. elegans*-*E. coli* system. Our data reveal that bacterial processing of glucose, rather than direct ingestion by the animal, results in reduced lifespan and healthspan, including reduced locomotion, oxidative stress resistance, and heat stress resistance in *C. elegans*. Chronic exposure of *E. coli* to glucose produces growth defects and increased advanced glycation end products within the *E. coli.* These negative effects are abrogated when the *E. coli* is not able to process the additional glucose and by the addition of the anti-glycation compound carnosine. Physiological changes of the host *C. elegans* are accompanied by dysregulation of detoxifying genes including glyoxalase, glutathione-S-transferase, and superoxide dismutase. Loss of *gst-4* shortens *C. elegans* lifespan and blunts the animal’s response to a glucose-fed bacteria diet. Taken together, we reveal that added dietary sugar may alter intestinal microbial *E. coli* to decrease lifespan and healthspan of the host and define the critical role of detoxification genes in maintaining health during a chronic high-sugar diet.

## Introduction

Approximately 0.1-5 % of the human microbiome consists of the bacterium *E. coli*, which presents little to no harm to its hosts (reviewed in ^1,2^). Despite this small fraction, *E. coli* is responsible for thousands of illnesses each year due to its rapid colonization of the human gut ^2,3^. Overall, the microbiota of the intestine plays a major role in human health and disease and undergoes large changes as we age ^4,5^. Additionally, recent studies suggest that a high sugar diet can alter the gut microbiota leading to age-associated illness ^6,7–9^. Direct evidence of the importance of the human intestinal microbiota and mechanistic analysis has not been possible due to its complexity, containing hundreds of types of organisms. In addition, there is very limited opportunity to perform experiments and as such, human microbiome studies are defined primarily by fecal samples which mainly represent only a small portion of the large intestine.

*C. elegans* present a highly malleable proxy to determine how diet affects microbes and the repercussions imparted onto their host. In the laboratory, the *C. elegans* diet consist of a single strain of *E. coli*, which then inhabits its intestine in a commensal relationship. As bacterivores, *C. elegans* have an obligatory symbiotic relationship with microbes as their food source that provide dietary nutritional supplementation ^10,11^. Recent studies have shown that the *C. elegans* bacterial diet can be altered to cause changes in lifespan and healthspan ^12,13,14^. Here, we have developed a *C. elegans*-*E. coli* system that allows direct modification of both the diet (*E. coli*) and the host (*C. elegans*) in response to changes in the environment. This simplified system takes advantage of *C. elegans* as a premiere system for studies on the aging process. Moreover, our studies highlight the effects of altering the health of *E. coli*, a significant component of the human intestinal microbiota.

Previously, studies on the effects of a high-glucose diet in *C. elegans* involved adding glucose either to the agar growth media directly or to the top of the agar media growth plate ^15–20^. This high glucose diet led to a decreased lifespan as well as reduced healthspan (locomotion), and changes in fat storage. In these studies, the additional glucose was in contact with both the bacteria as well as *C. elegans*, therefore we questioned the source of the high glucose effect. Many of the protocols have variations with regard to how the glucose was applied to the agar, whether the bacteria were alive or dead, how the bacteria were killed, and whether the bacteria had any direct contact with the glucose ^15–20^. To separate the effects of *C. elegans* consuming a glucose fed bacteria diet, we developed a new experimental procedure based on previous studies from ^21^. In our protocol, prior to seeding the bacteria on the plate, it is incubated with glucose for three days.

Added dietary sugar has been associated with an alarming rise in a multitude of debilitating medical conditions including obesity, diabetes, cardiovascular and neurodegenerative diseases, such as Alzheimer’s disease, due to its impact on the generation of advanced glycation end products (AGEs). Both exogenous and endogenous sources contribute to the levels of AGEs, a group of heterogeneous compounds, mainly proteins and lipids, that become glycated as a result of exposure to sugars. Glycation results from a non-enzymatic reaction where the carbonyl group of reducing sugars is covalently coupled to proteins, lipids and nucleic acids. Endogenous formation of AGEs occurs continuously at low levels, but exogenous sources, including consumption of a high-sugar diet, can drastically increase this pool. As humans age and in certain diseased states, AGE levels rise and are associated with increased cardiovascular risk, diabetes, chronic kidney disease, and Alzheimer’s disease.

Many studies in animal model systems show that dietary consumption of AGEs contributes to oxidative stress and inflammation which could contribute to a number of chronic disease states (reviewed in ^22,23^). Additionally, dietary AGEs have been shown to affect the inflammatory response in cell culture ^24^, but there are mixed results for the effects of dietary AGEs in human trials ^22^. Emerging studies also suggest that dietary AGEs can contribute to the onset of organ damage, affect metabolic control, and therefore impact global health ^23^.

We find that our *C. elegans*-*E. coli* system, where we modify the health of the microbiome by modulating dietary sugar, results in changes in lifespan and healthspan of *C. elegans*. Increased added dietary sugar results in decreased lifespan, decreased healthspan including decreased movement in liquid, and decreased oxidative stress resistance, only when the *E. coli* have been continuously subjected to additional dietary sugar. Addition of the anti-glycation compound carnosine ameliorates the negative effects of glucose in the diet. Added dietary sugar suppresses *C. elegans* oxidative stress resistance, most notably by suppression of *gst-4* expression. Our data reveal a central role for *gst-4* in the regulation of a high-sugar diet. Both a glucose-fed bacterial diet as well as loss of *gst-4* shorten lifespan. In addition, loss of *gst-4* blunts the animals’ response to a glucose-fed bacteria diet. Taken together, our model system allows dissection of a microbiota on a level not possible in humans, with univariable analysis of both the microbiota and the host which reveals the intimate connection between oxidative stress and a high-sugar diet.

## Results

Previous studies on the effects of glucose toxicity in *C. elegans* involved several variables, including whether the bacterial diet was alive or dead, and whether the additional glucose was added either to the molten agar or to the top of the agar growth plate ^15–20^. Therefore, the additional glucose was in contact with both the bacteria as well as *C. elegans*.

To separate the effects of *C. elegans* consuming a bacteria diet with added glucose, we developed a new experimental procedure based on studies from our previous bacterial studies^21^. In our system, OP50 *E. coli* were inoculated in LB media supplemented with various glucose concentrations (ranging from 0-0.8%) for 3 days; followed by heat killing of the *E. coli* which is then seeded onto nematode growth medium (NGM) plates for physiological and biochemical assays (Figure 1a). Heat killing of the bacterial diet prevented any possible bacterial proliferation differences. This 3-day glucose exposure resulted in the OP50 *E. coli* exhibiting a significant decrease in colony forming units (Figure 1b). The optical density of the glucose-fed *E. coli* was not significantly changed, indicating that *C. elegans* were consuming similar biomass amounts (Supplementary Figure S1a). We also confirmed the glucose-fed *E. coli* had elevated levels of intracellular glucose (Supplementary Figure S1b). Similar to our previous studies with a different strain of *E. coli* supplemented with glucose^21^, the 3 day incubation of OP50 *E. coli* supplemented with glucose also led to an increase in the AGE carboxymethyl-lysine (CML) as detected by ELISA (Figure 1c). Therefore, the *E. coli* shown in Figure 1b grown with supplemental 0.4% glucose show an increase in CML along with the loss of CFU. Together, from our initial studies, we selected the 3 day period of growth with supplemental 0.4% glucose for our experimental paradigm.

**Figure 1.**
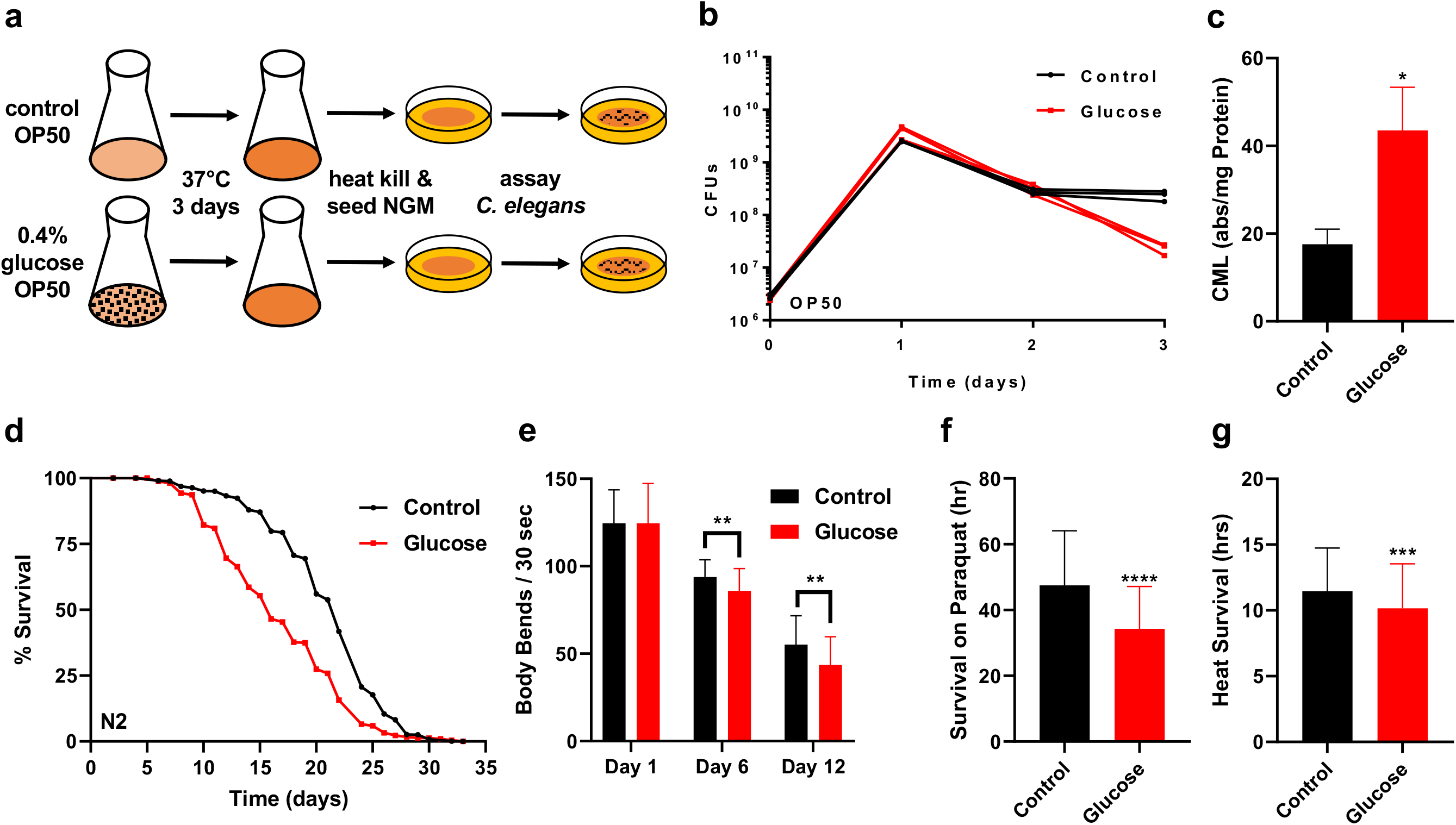
Effects of glucose fed bacteria. (A) Experimental setup for OP50 *E. coli* processing of glucose and assaying of *C. elegans* on the diet. (B) Effect of 0.4% glucose supplementation on OP50 *E. coli* colony forming units over time. (C) CML Assay performed on control and 0.4% glucose fed OP50 *E. coli*. (D) Lifespan assay of wild type *C. elegans* treated with control and 0.4% glucose fed OP50 *E. coli*. (E) Movement/swimming ability of *C. elegans* treated with control and 0.4% glucose fed *E. coli* over time. (F) Oxidative stress resistance of wild type *C. elegans* treated with control and 0.4% glucose fed *E. coli* for 6 days, measured by survival on paraquat media. (G) Heat stress resistance of wild type *C. elegans* treated with control and 0.4% glucose fed *E. coli* for 6 days, measured by survival at 37°C.

*C. elegans* consuming the glucose-fed *E. coli* diet show a significant reduction (~30%) in lifespan compared to animals fed a control diet with no added glucose (Figure 1d). Importantly, the reduced lifespan was also accompanied by a reduction in healthspan. As shown in Figure 1e, animals consuming the glucose fed bacteria diet, show a reduction in healthspan as quantified by movement in liquid/swimming/thrashing. To further evaluate health parameters, oxidative and heat stress resistance were tested on *C. elegans* consuming the glucose fed *E. coli* diet for 6 days (Figures 1f-g). We selected day 6 as the time point for further analysis as these reflected differences that could be observed prior to any effects on *C. elegans* viability. An additional phenotype of the glucose fed bacteria diet is that animals show a consistent reduction in size over time (Supplementary Figure S1c). Therefore, our results illustrate that consumption of the glucose-fed bacteria diet mirrors results seen in more traditional glucose supplementation experiments ^15–20^. Together, these data reveal that *C. elegans* consuming a diet of glucose-fed *E. coli* show significant reduction in lifespan and reduced healthspan.

Next, we examined how the bacterial processing of glucose affected the health of *C. elegans* using three different methods. First, we supplemented *E. coli* with glucose prior to the culture start (Pre glucose; method as in Figure 1a) compared to supplementation after 3 days and heat kill (Post glucose). In the latter method (Post glucose), glucose was readily available only to *C. elegans*, in comparison to the former diet, in which glucose was incubated with the *E. coli* first. As shown in Figure 2a, lifespan was shortened only when the live *E. coli* diet was allowed to process the glucose. Remarkably, although the post glucose supplementation had a higher amount of glucose (80×) available to the animal (Figure 2g), consumption of the post-glucose diet showed no change in lifespan (Figure 2a). In addition to the effect on lifespan, oxidative stress resistance was reduced only in the pre-glucose diet (Figure 2d). Movement in liquid required a greater time to see the effects of the consumption of the glucose diet since there were no significantly changes until day 12 (Supplementary Figure S2b). Therefore, bacterial processing of glucose decreases lifespan and oxidative stress resistance. Notably, prior studies in mammals have already linked oxidative stress to consumption of a high AGE diet ^23,25–27^.

**Figure 2.**
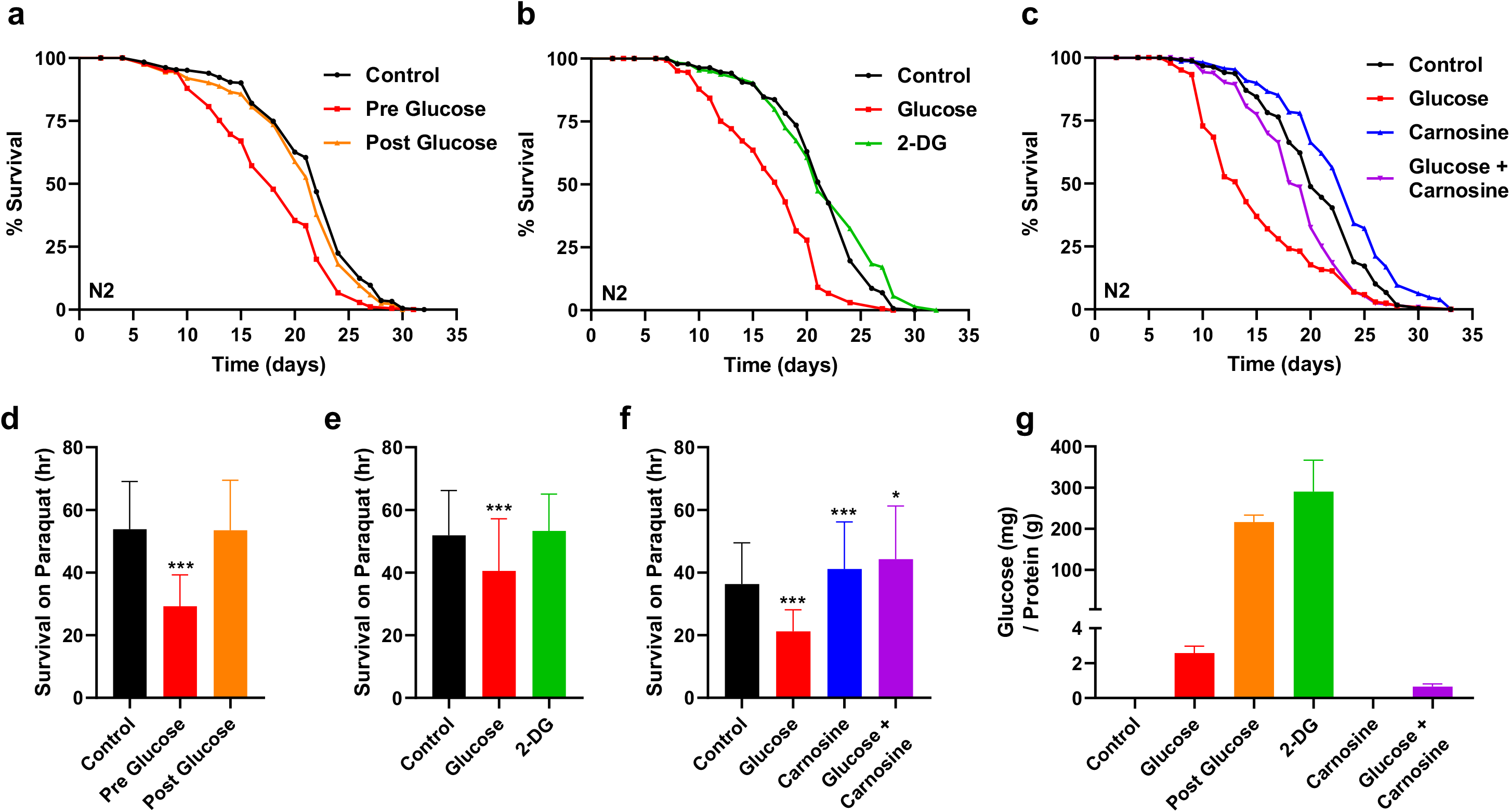
Mitigating bacterial metabolism of glucose alters its effects on *C. elegans*. (A) Lifespan assay of wild type *C. elegans* treated with control and *E. coli* that was supplemented with 0.4% glucose pre culture or post culture. (B) Lifespan assay of wild type *C. elegans* treated with 0.4% glucose or 0.4% 2-deoxy-glucose (2-DG) fed OP50 *E. coli*. (C) Lifespan of wild type *C. elegans* treated with control, 0.4% glucose, 50mM carnosine, and 0.4% glucose + 50mM carnosine fed OP50 *E. coli*. (D) Oxidative stress resistance of wild type *C. elegans* treated with control and *E. coli* that was supplemented with 0.4% glucose pre culture or post culture, measured by survival on paraquat media. (E) Oxidative stress resistance of wild type *C. elegans* treated with 0.4% glucose or 0.4% 2-deoxy-glucose (2-DG) fed OP50 *E. coli*. measured by survival on paraquat media. (F) Oxidative stress resistance of wild type *C. elegans* treated with control, 0.4% glucose, 50mM carnosine, and 0.4% glucose + 50mM carnosine fed OP50 *E. coli*, measured by survival on paraquat media. (G) Glucose assay performed on *E. coli* that was supplemented with control, 0.4% glucose pre culture or post culture, 0.4% 2-deoxy-glucose (2-DG), 50mM carnosine, and 0.4% glucose + 50mM carnosine fed OP50 *E. coli*, results normalized to protein concentration.

A second method to examine the effects of bacterial metabolism of glucose on the *C. elegans* host was the use of the synthetic glucose analog 2-deoxy-D-glucose (2-DG). When consumed, 2-DG is phosphorylated by hexokinase which then cannot be further processed and therefore is used as a glycolytic inhibitor. We treated the bacterial diet with 2-DG. *C. elegans* consuming the bacterial diet with 2-DG have a normal lifespan accompanied by no change in either resistance to oxidative stress or movement in liquid (Figures 2b, 2e, and Supplementary Figure S2b). Thus, consuming the bacterial diet with 2-DG results in phenotypes opposite to those seen with a glucose fed bacteria diet.

Third, to further elucidate the connection between microbial processing of glucose and host health, we used the dipeptide carnosine in combination with glucose. We have previously shown that carnosine has anti-glycation properties that reduce the toxic effects of glucose in *E. coli* ^21^. As shown in Figure 2c, as expected, carnosine fed *E. coli* significantly extends *C. elegans* lifespan and ameliorates the effect of added glucose. Interestingly, there is little to no effect of carnosine on *C. elegans* when the compound is mixed within the agar of the plate then seeded with heat-killed *E. coli* (Supplementary Figure S2c). Additionally, glucose concentrations within the *E. coli* are decreased by 3x when the *E. coli* are fed both glucose and carnosine when compared to glucose alone (Figure 2g). *C. elegans* consuming this diet are protected from the oxidative stress resistance decrease that occurs in glucose supplementation (Figure 2f). Therefore, across three different methods, we reveal that bacterial processing of glucose dictates *C. elegans* lifespan and healthspan.

Animals consuming the glucose fed bacteria exhibit a shortened lifespan and decreased stress resistance. Therefore, we next examined whether the expression levels of key genes involving environmental defenses could account for the observed phenotypes from the glucose fed bacteria diet using both transgenic GFP animals as well as RT-qPCR. We used known stress-related transgenic GFP strains to observe any potential changes in the dynamics of expression associated with consumption of glucose-fed bacteria within the host over time. To determine the downstream signaling pathway in the host that responds to the glucose fed bacteria diet, we screened distinct pathways induced by oxidative stress *(gst-4, sod-3)*, or heat stress *(hsp-16.2)*, the mitochondrial UPR *(hsp-6)*^28,29^, the glyoxalase system *(glod-4, gcs-1)*, the transcription factor *skn-1* ^30^, and the insulin/IGF-1 signaling pathway FOXO, *daf-16* (Figure 3a and Supplementary Figure S3a).

**Figure 3.**
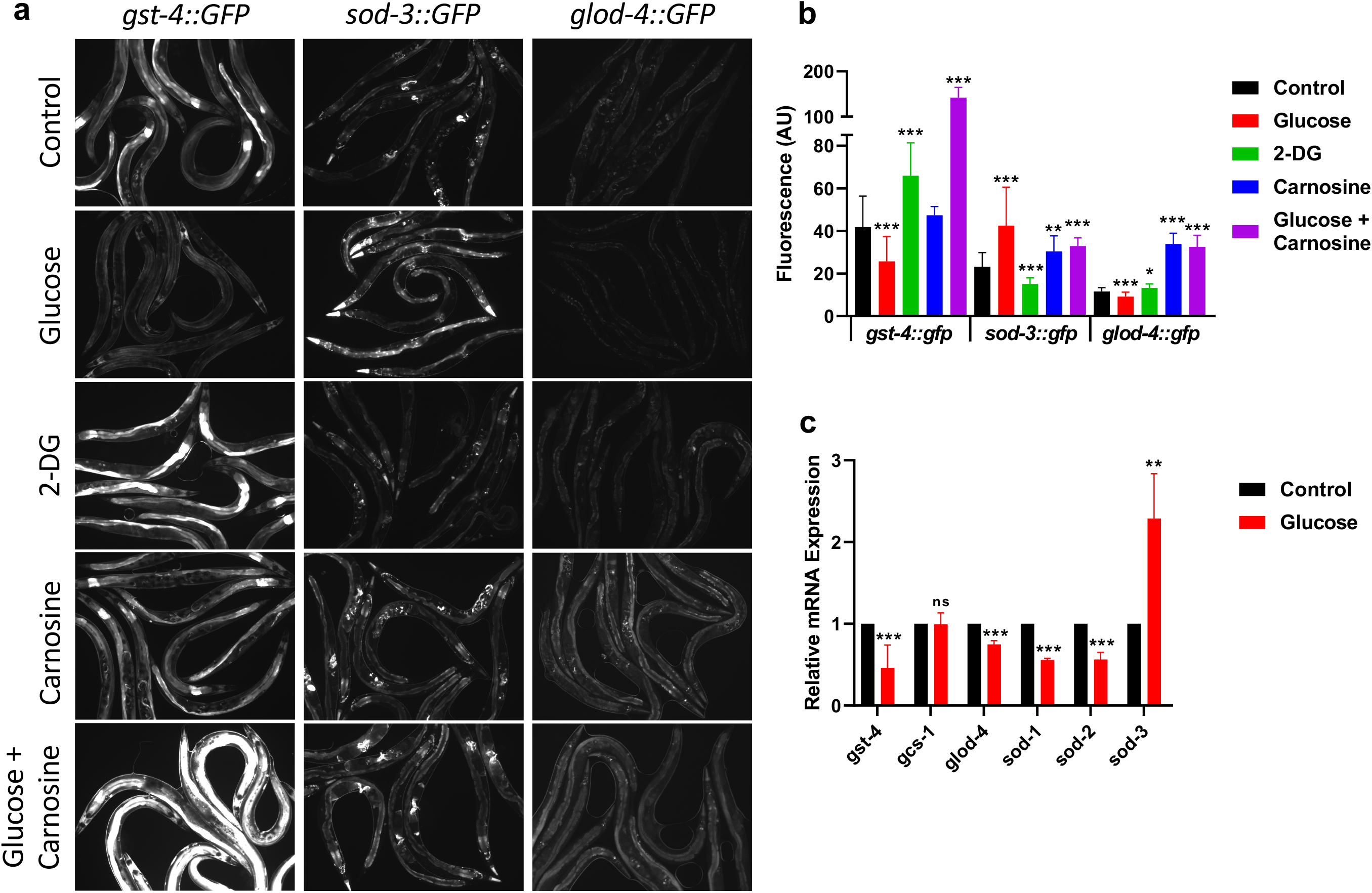
Bacterial metabolism of glucose on host gene expression. (A) Fluorescent imaging of transgenic strain *C. elegans* [*gst-4∷gfp*], [*sod-3∷gfp*], or [*glod-4∷gfp*], after treatment with control, 0.4% glucose, 0.4% 2-deoxy-glucose (2-DG), 50mM carnosine, and 0.4% glucose + 50mM carnosine fed OP50 *E. coli* for 6 days. (B) Fluorescence quantification of transgenic strain *C. elegans* [*gst-4∷gfp*], [*sod-3∷gfp*], or [*glod-4∷gfp*], after treatment with 0.4% glucose, 0.4% 2-deoxy-glucose (2-DG), 50mM carnosine, and 0.4% glucose + 50mM carnosine fed OP50 *E. coli* for 6 days. (C) RTqPCR experiments performed using wild type *C. elegans* treated with control or 0.4% glucose fed *E. coli* for 6 days.

Notably, as shown in Figure 3a, we observed blunted expression of *gst-4∷GFP* and *glod-4∷GFP* when animals consumed the glucose-fed bacteria diet, as well as an increase in *sod-3∷GFP*. To remove any bias associated with the visual assay, we quantified the fluorescence of those animals (Figure 3b). Then, to confirm the validity of our findings, we performed RT-qPCR on wild-type animals under the same conditions. We find consistent gene expression changes when assayed by RT-qPCR (Figure 3c).

Next, we tested if gene expression changes paralleled our findings in Figure 2 with the use of the glucose analog 2-DG, which did not alter *C. elegans* lifespan or healthspan. As shown in Figure 3a, we observed gene expression levels opposite to glucose when animals were fed a bacterial diet supplemented with 2-DG. This finding was confirmed with fluorescence quantification (Figure 3b). Therefore, in response to a high glucose diet where bacteria process the glucose, we observed a decrease in lifespan and healthspan associated with a downregulation of several key detoxification genes.

Then, we checked GFP expression changes in *C. elegans* when fed a diet consisting of *E. coli* fed carnosine alone or in combination with glucose as carnosine ameliorated the shorter lifespan induced by glucose (Figure 2). *C. elegans* consuming carnosine-fed *E. coli* exhibit a mild increase in lifespan and also show a slightly higher level of *gst-4∷GFP*, *sod-3∷GFP*, and *glod-4∷GFP* expression (Figures 3a and 3b). When carnosine is added to the glucose-fed *E. coli*, lifespan is greater than glucose alone and *gst-4∷GFP* fluoresces significantly brighter. *sod-3∷GFP* and *glod-4∷GFP* show similar expression to carnosine alone (Figures 3a and 3b). Furthermore, *C. elegans* consuming carnosine-fed *E. coli* are protected from the oxidative stress resistance decline that occurs during glucose supplementation and show a stabilization of gene expression as part of the stress response (Figures 2f and 3a).

We also screened two additional signaling pathways: *skn-1* (Nrf2) and *daf-16* (insulin/IGF-1). As shown in Supplementary Figure S3a, the glucose fed-bacteria diet did not change expression of *skn-1op∷GFP* nuclear translocation. Since *skn-1* showed no changes in GFP expression in response to the glucose-fed bacteria diet, we further tested one of its target genes, *gcs-1*, often used as a surrogate for *skn-1*. There was neither change in *gcs-1* mRNA levels (Figure 3c), nor *gcs-1∷GFP* expression (Supplementary Figure S3a). We also examined the insulin/IGF signaling pathway in response to the glucose fed bacteria diet by nuclear translocation of *daf-16a∷GFP.* We did not observe any changes in nuclear translocation for *daf-16a∷GFP*. Consistently, RT-qPCR showed little change for both *skn-1* and *daf-16* (Supplementary Figure S3c). Therefore, taken together, our results illustrate that in response to the glucose fed bacteria diet, the glycated bacteria consistently promote an environment of oxidative stress similar to consumption of a high AGE diet in mammals ^23,25–27^.

Surprisingly, we observed a consistent upregulation of *sod-3* and *sod-5*, which presumably should increase resistance to oxidative damage^31^. A recent report by Dues et al. (2017) ^32^ examined the total superoxide dismutase capacity of *C. elegans*. They found that *sod-3* and *sod-5* together only account for 1.7% of all *sod* expression while *sod-1*, *sod-2*, and *sod-4* together amass 98.2% of the total *sod* mRNA. In accordance with these findings, we mathematically extrapolated our RTqPCR results onto the published expression ratios to illustrate total *sod* expression (Figure 3c and Supplementary Figure S3b). Animals consuming glucose fed bacteria show a ~50% decrease in *sod-1* and *sod-2* expression which overshadows any increase in *sod-3* expression, and therefore overall, the glucose-fed bacteria-treated animals exhibit a reduction of *sod* expression to only 60% of total wild type *sod* mRNA capacity (Supplementary Figure S3b).

Therefore, across multiple experimental paradigms, our data confirm that bacterial processing of glucose promotes a decrease in lifespan as well as a reduction in health, and stress resistance. These phenotypical changes are accompanied by differential regulation of glyoxalase, glutathione-S-transferases, and superoxide dismutase - all of which should enzymatically detoxify the glycolytic effects of glucose.

Our data reveal that glucose-fed bacteria which are high in AGEs promote an environment that causes animals to exhibit signs of oxidative stress including strong and consistent suppression of *gst-4* (glutathione transferase-4) involved in the regulation of the Phase II oxidative stress response ^33^. Analysis of transcriptional activation of *gst-4* is often used as a read-out for oxidative stress tolerance ^34^. Therefore, we next generated a loss of function allele of *gst-4* by CRISPR/Cas9.

We generated two independent isolates of the *gst-4* mutation, *lp10* and *lp11* by CRISPR/Cas9. As shown in Figure 4a, sequencing *lp11* revealed a loss of 599 bp deletion than spans all 4 exons of wild type *gst-4*. Next, we tested the *gst-4* mutants for lifespan and show that the *gst-4* mutation results in a decrease in lifespan (Figure 4b). We further examined whether loss of *gst-4* interfered with the high sugar diet. As shown in Figure 4c, mutation in *gst-4* abrogates the response to the glucose fed bacteria diet. Therefore, our results indicate that *gst-4* is necessary to respond to a high glucose fed bacterial diet. Together, our data reveal that dietary AGEs promote oxidative stress in the host dependent on the Phase II oxidative stress response.

**Figure 4.**
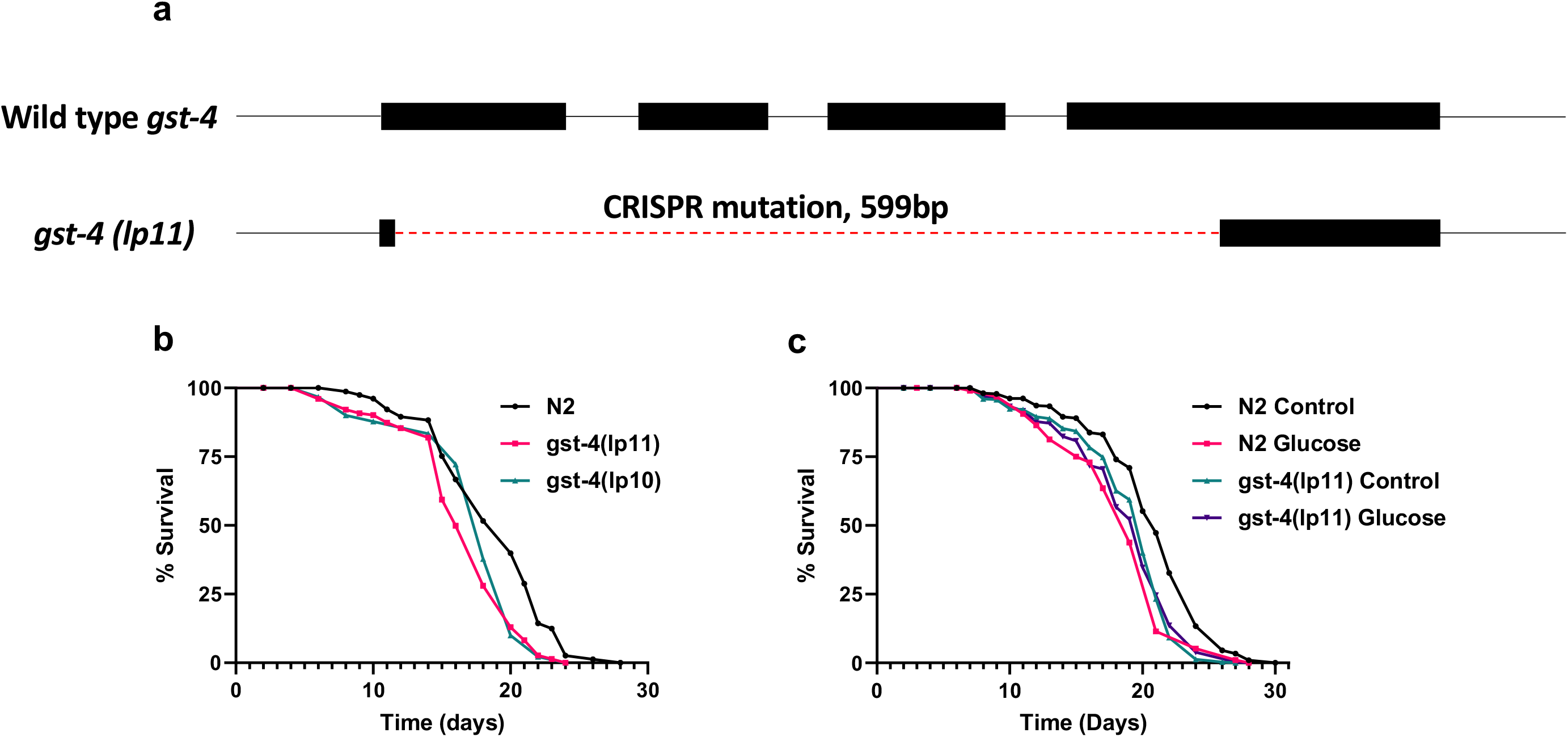
Generation and characterization of a *gst-4* mutant with its response to glucose fed *E. coli*. (A) Diagram of genomic wild type *C. elegans gst-4* DNA sequence and the *gst-4(lp11)* allele DNA sequence, lines represent introns and non-coding regions, rectangles represent exons, and the dotted line represents the *lp11* allele CRISPR mutation. (B) Lifespan assay of wild type N2, *gst-4(lp10)*, and *gst-4(lp11)* mutant *C. elegans* on OP50 NGM. (C) Lifespan assay of wild type N2 and *gst-4(lp11)* mutant *C. elegans* treated with control and 0.4% glucose fed OP50 *E. coli*.

## Discussion

As bacterivores, the diet of *C. elegans* in the laboratory typically consists of a single *E. coli* strain, which then inhabits its gut in a commensal relationship. Here, we test the *C. elegans-E. coli* interaction in response to changes in the dietary environment. Previous studies examining a high sugar diet involved several different methods. A given percentage of glucose, typically 2%, is either added to the agar before cooling or added to the top of the agar filled plate after it has cooled. Bacteria, the diet for *C. elegans*, would then be spread onto the agar plate. In such an experimental paradigm, there is a three way interaction: *E.coli* is exposed to the glucose, *C. elegans* is exposed to the glucose, and *C. elegans* consumes the *E.coli* exposed to the glucose. ^15–20,35–37^. In this case, as shown in our model in Supplementary Figure S1d, glucose has both direct and indirect contact with *C. elegans* and there is no differentiation between the direct glucose effect onto *C. elegans*, or the indirect effects of glucose on the *E. coli.* Accordingly, some studies have opted to utilize inactivated *E. coli* to examine direct effects onto *C. elegans*.

In contrast here, our *C. elegans*-*E. coli* system reduces the variable differences and specifically examines bacterial glucose processing and its effect on *C. elegans*. Addition of glucose to the bacterial culture allows processing by the bacteria to occur preceding any exposure to *C. elegans*. These glucose-fed bacteria are then seeded onto the plate (the diet), which is the indirect bacterial glucose effect on *C. elegans.* In our indirect method, we find that *C. elegans* consuming the glucose-fed *E. coli* diet have a 24% reduction in lifespan and a reduction in healthspan including locomotion and oxidative stress resistance (Figure 1, Supplementary Table S1). Therefore, both the indirect and direct effects of glucose result in poor healthy aging ^15–20^.

Our data reveal that the specific bacterial processing of glucose negatively affected the health of *C. elegans* by using three different methods to separate, inhibit, and suppress bacterial processing (Figure 2). First, we supplemented the *E. coli* culture with the same concentration of glucose, but addition was after the bacterial diet was heat-inactivated. This direct application of glucose had no effect on lifespan. This may be due to the fact that the amount of glucose added to the top of the plate was relatively compared to previous reports utilizing 2% per plate. However, this dose, when processed by bacteria, drastically changes the health of the host. Secondly, we incubated the *E. coli* with glucose analog 2-DG to inhibit glycolysis within the bacteria. In contrast to glucose consumption, 2-DG results in healthier animals with a normal lifespan. In a third set of experiments, we incubated the *E. coli* culture with carnosine. We show that with glucose supplementation, the *E. coli* increase their levels of AGEs by 3-fold. Carnosine, an anti-glycation compound, has the ability to make glucose less toxic as it can reduce AGEs ^21^. We observed that carnosine alone as well as carnosine in combination with glucose resulted in healthier animals when on a glucose-fed bacteria diet. Therefore, across three methods, our data reveal that the bacterial processing of glucose dictates the phenotypes of the animals consuming this diet.

Consistently, the glucose fed *E. coli* shortened the median lifespan by ~35% (Supplementary Table S1). Further, when fed the glucose and carnosine treated *E. coli*, lifespan reduction was reduced only by ~7%. Carnosine alone treated *E. coli* increased lifespan by ~7%. Additionally, when applied directly to the diet, carnosine supplementation had little to no effect on *C. elegans*. Thus, benefits were only observed when carnosine was processed by the bacteria. Carnosine treated *E. coli* also increased *C. elegans* healthspan as measured by resistance to oxidative stress, both alone and in the presence of glucose. Together, these results suggest that carnosine used as a prebiotic intervention may have the potential to affect the resident microbiota and alleviate dietary glucose related oxidative stress.

Previous studies have shown a strong connection between AGEs and oxidative stress ^23,38,39^. We show that when animals consume the highly glycated diet caused by the additional sugar, they are subject to high levels of oxidative stress. We surveyed a broad panel of stress reporter genes when animals are subject to the glucose fed bacteria diet (Figure 3). Animals consuming the glucose-fed bacteria diet exhibit a down regulation of *gst-4*, both at the GFP expression as well as the mRNA levels. Interestingly, our data show that the microbial processing of glucose inhibits the expression of *gst-4*. Further, our analyses reveal that the glucose fed bacteria diet promotes an environment of constant oxidative stress such that when *C. elegans* age on this diet, they become more susceptible to oxidative stress. Furthermore, our data reveal that loss of *gst-4* prevents the animal from responding to the high-sugar/glycated diet (Figure 4). It should be noted that we examined multiple other glutathione-S-transferases including *gst-10, gst-29* and found that they were all downregulated. This indicates the possibility that the bacterial processing of glucose suppresses the animal’s ability to respond to this oxidant rich environment, which may perpetually cause chronic vulnerability to oxidants.

We suggest that using our *C. elegans*-*E. coli* system, it is possible that we mimic dietary effects on bacteria within the human digestive tract. We use a long-term bacterial culture with limited resources and analyze the effect on its host. We suggest this is similar to the entrenched bacteria within the human digestive tract. Interestingly, Crohn’s disease is characterized by bacterially derived persistent inflammation of the intestine and colon and therefore could be the physiological manifestation of this condition. In fact, recent clinical studies find that patients suffering from Crohn’s Disease/IBD show oxidative stress ^40^. Therefore, the blunted transcriptional response from consumption of the glucose-fed bacteria could help in future work on the dynamics involved in persistent bacterial inflammation in cases of Crohn’s disease.

Taken together, our data show that bacterial processing of glucose resulted in reduced lifespan and reduced stress resistance in their host with downregulation of stress response genes (Figure 5). Transcriptional response by *C. elegans* to their nutritional intake is protected by the preventative mechanisms of action of carnosine. Consumption of the high glucose/highly glycated/high AGEs diet shortens lifespan, reduces healthspan and promotes oxidative stress.

**Figure 5.**
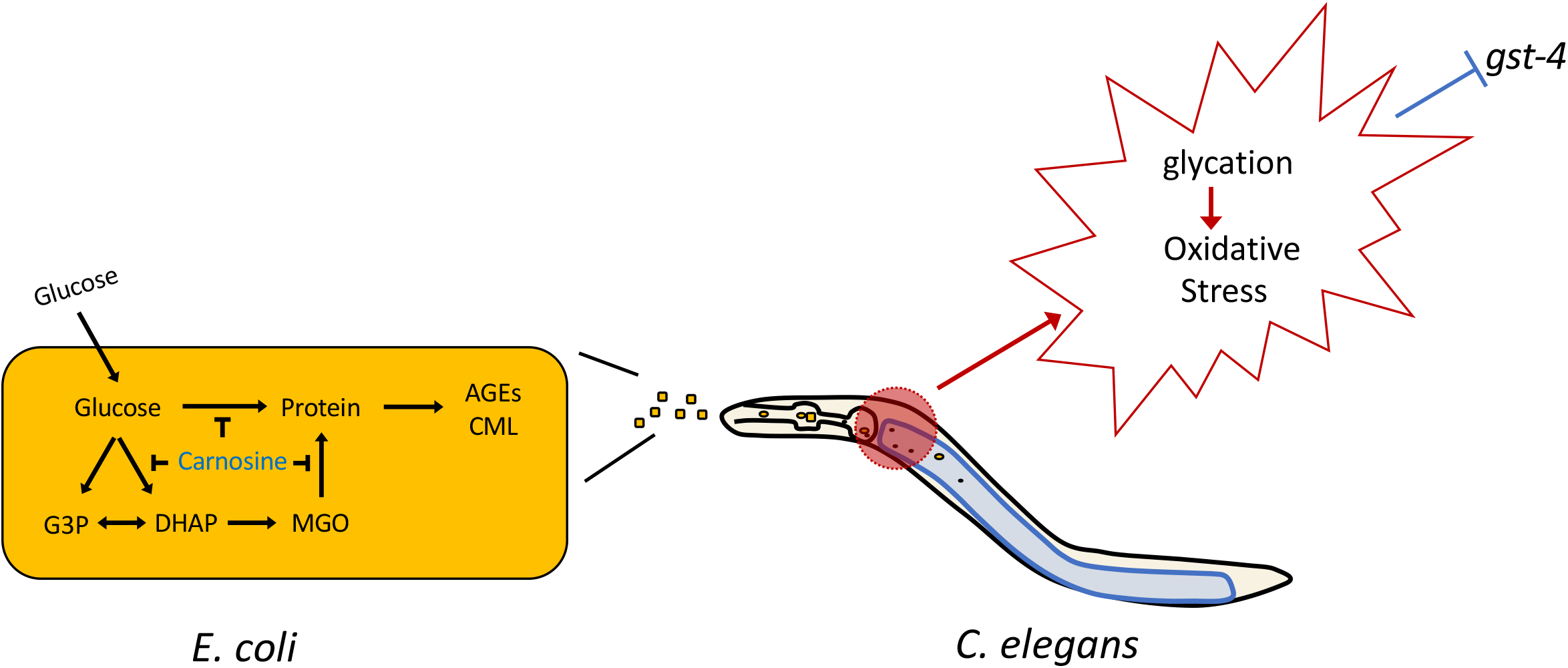
Model of *E. coli* glucose processing effects onto *C. elegans*.

## Methods

### Strains

All strains were maintained at 16°C using standard *C. elegans* techniques except where indicated 41. Strains used in this study: N2, *daf-16(mgDf50)* I, *daf-2(e1370)* III, TJ356 [*daf-16∷gfp*], LD1008[ldEx9 [*skn-1(operon)∷GFP + rol-6(su1006)*]], BC15643 [*dpy-5(e907)* I; sEx15643[*rCes C16C10.10∷GFP + pCeh361*]], CL2166 [dvIs19 [*(pAF15)gst-4p∷GFP∷NLS*] III.], CF1553 [muIs84 [*sod-3p∷GFP*]], SJ4100 [zcIs13[*hsp-6∷GFP*]], CL2070 [dvIs70[*hsp-16.2p∷GFP + rol-6(su1006)*]], LD1171 [ldIs3 [*gcs-1p∷GFP + rol-6(su1006)*]], Some strains were provided by the CGC, which is funded by NIH Office of Research Infrastructure Programs (P40 OD01044).

### Lifespan Assay

All lifespan assays were performed at 20°C. Strains were semi-synchronized by allowing gravid adults to lay eggs on standard NGM plates overnight, and animals developed for several days until they reached L4. Then, ∼100 L4s were transferred to NGM or treatment plates and kept at 20°(∼33 animals per plate). Animals were scored by gently tapping with a platinum wire every 2-3 days. Animals that did not respond were scored as dead. Animals that crawled off the plate, bagged, or died from vulva bursting were censored from the analysis. Each figure shows data collected from all trials. The mean lifespan, SD, and P values were calculated via two-tailed unpaired t tests (Supplementary Table S1). Survival graphs and statistical analyses were produced using GraphPad Prism 8 (GraphPad Software).

### Glucose/Carnosine fed *E. coli* Diet

Standard Nematode Growth Media (NGM) ^41^ was autoclaved at 121°C for 25 min, then cooled at room temperature (RT) and 100μg/mL Ampicillin (Fisher Scientific) was added prior to pouring plates. Plates dried for 3 days at RT before seeding with *E. coli*. The OP50 *E. coli* was grown for 3 days in LB only (control), or LB supplemented either pre-culture or post-culture with 0.4% D-(+)-glucose (Sigma), 0.4% 2-Deoxy-D-glucose (Sigma), and 50mM L-Carnosine (Sigma). After 3 days, the *E. coli* cultures were subjected to a 65°C heat bath for 30 min to inhibit further growth. OP50 was then streaked onto an LB agar plate and cultured at 37°C overnight to confirm lack of growth. Plates were stored at 4°C until use.

### Glucose Assay

OP50 *E. coli* was grown for 3 days in LB only (control), or LB supplemented either pre-culture or post-culture with 0.4% D-(+)-glucose (Sigma), 0.4% 2-Deoxy-D-glucose (Sigma), and 50mM L-Carnosine (Sigma). After 3 days, the *E. coli* cultures were subjected to a 65°C heat bath for 30 min to inhibit further growth. Samples were diluted in reaction buffer to accurately detect glucose levels and frozen at −20°C until use. Glucose amount was determined using the Amplex Red Glucose/Glucose Oxidase Assay Kit (Invitrogen), in triplicate. Protein concentration was then measured with Pierce™ Coomassie Plus (Bradford) Assay Kit (Thermo Scientific). Glucose levels were standardized to protein levels, and then used to graph over three replicate experiments using GraphPad Prism 8 (GraphPad Software).

### RNA Extraction and RT-qPCR

Approximately 200 synchronized L4 stage animals were transferred to treatment plates and grown at 20°C for 6 days, then washed off the plates with M9 buffer and rinsed twice with DEPC-treated water. Total RNA was isolated using TRIzol Reagent with the Direct-zol RNA MiniPrep (Zymo Research). After RNA extraction, first-strand cDNA was synthesized from 1.0 μg of total RNA using dNTPs, Oligo(dT)12–18 and SuperScript II Reverse Transcriptase (Invitrogen). Quantitative PCR was done with an Applied Biosystems StepOne Plus Real-Time PCR system with Power SYBR Green PCR Master Mix (Applied Biosystems) per the manufacturer’s instructions, with triplicates done for each of three biological replicates, and *act-1* used as the endogenous control for relative expression normalization. Sequences of primers can be found in Supplementary Table S2. Specificity of PCR amplification was determined by the melting curve for each reaction. The threshold cycle (CT) for each primer set was automatically determined by the StepOne Software 2.1 (Applied Biosystems). Relative fold changes of gene expression were calculated by using the 2^−^△△Ct method. The RQ values were then used to graph relative gene expression over at least three replicate experiments using GraphPad Prism 8 (GraphPad Software).

### Healthspan: Body Bends/Movement in Liquid Media

Synchronized L4 stage animals were transferred to treatment plates and incubated at 20°C for 1, 3, 6, or 12 days. On the experimental day, individual animals were picked onto an unseeded NGM plates, then 30μL of M9 buffer was pipetted onto each animal. After 5 seconds, the number of body bends was recorded over 30 seconds using a mounted DMK 21AF04 camera (The Imaging Source) outfitted onto a dissecting microscope. At least 20 animals were recorded per strain or treatment and the average body bends per minute for each of the samples was calculated and graphed using GraphPad Prism 8 (GraphPad Software).

### Healthspan: Resistance to Oxidative Stress

Synchronized L4 stage animals were transferred to treatment plates and incubated at 20°C for 6 days. Animals were then transferred to paraquat plates at 20°C and scored twice daily for survival. Animals were touched with a platinum wire, and those that did not respond were scored as dead. Two independent replicates were performed, and the mean survival was calculated using GraphPad Prism 8 (GraphPad Software). Paraquat plates were made by adding 1mL of 250mM paraquat solution to corresponding treatment plates. Then plates were put on a shaker for 1 hour, followed by 1.5 hours in a laminar flow hood to ensure plates were dry with the paraquat evenly distributed.

### Resistance to Heat Stress

Synchronized L4 stage animals were transferred to treatment plates and incubated at 20°C for 6 days. Animals were transferred to 37°C and then scored every 2 hours for survival. Animals were touched with a platinum wire, and those that did not respond were scored as dead. Two independent replicates were performed, and the mean survival was calculated using GraphPad Prism 8 (GraphPad Software).

### Fluorescence Imaging/Quantification

Synchronized L4 stage animals were transferred to treatment plates and incubated at 20°C for 1, 3, 6, or 12 days. On the experimental day, 10-12 animals were then picked onto 2% agarose pad on a glass slide and immobilized with 100mM NaN3. Fluorescence imaging of GFP was done using a Zeiss Axioskop 2 plus fitted with a Hamamatsu ORCA-ER camera and a FITC filter. Images taken were in grayscale. Quantification of GFP expression was performed using ImageJ software. In the photos, each worm where at least 95% was in frame were outlined by hand and then measured for minimum, maximum, and average pixel intensity within the defined area. The minimum pixel intensity recorded was then subtracted to remove background fluorescence interference. The average pixel intensity per worm compiled across 2 or more batch experiments was then plotted using GraphPad Prism 8 (GraphPad Software).

### *E. coli* AGE assay

Performed an ELISA to detect carboxymethyl-lysine levels as in Pepper et al (2010)^21^ with 100 μl of supersensitive, 3,3’,5,5’-Tetramethylbenzidine (TMB) (Sigma) used as the color reagent and read at 630 nm on a BioTek ELx808 absorbance reader.

### CRISPR/Cas9 generation of a *C. elegans* mutant

Guide RNA sequences were designed using http://crispor.tefor.net ^41^ and genome editing was performed as described previously ^42,43^. Cas9, tracrRNA and two crRNAs targeting the coding region of *gst-4* were injected as ribonucleoprotein (RNP) complexes into gonads of N2 hermaphrodites. Using Prf4:*rol-6(su1006)* as injection marker, F1 Roller progeny were cloned and genotyped for deletions. Genotyping was performed with oligos flanking the guide binding sites to identify mutants that lack the entire region between the crRNA target sites. Genotyping PCR primer sequences were designed utilizing the aid of IDTdna PrimerQuest tool and were manufactured by Invitrogen. Two such deletion mutants were isolated as heterozygotes and later homozygosed to obtain *gst-4 (lp10)* and *gst-4(lp11)*.. After PCR, the sequence was confirmed with DNA sequencing performed by GENEWIZ, Inc.

### *E. coli* Fitness (Survival) Assay

Overnight cultures were inoculated from frozen stocks. 100μl of each strain was spread on each plate in triplicate and allowed to dry by evaporation. At each time point, the number of viable cells was determined by “coring” a section of the lawn with a sterile Pasteur pipet (inner diameter ~1.5mm), resuspending each core in 50 μl of LB, and titering by serial dilution and plating onto LB medium; the limit of detection is <100 cfu/ml ^45,46^.

## Acknowledgements

We are grateful to members of the Tissenbaum and Finkel lab for advice and suggestions, Susan Lee and Evelyn Caez for technical support, Dr. Craig Mello for advice and support with generation of the *gst-4* mutant, and Dr. Jeremy Van Raamsdonk for advice and primer sequences. Some of the *C. elegans* strains were kindly provided by the *Caenorhabditis* Genetics Center, which is funded by NIH Office of Research Infrastructure Programs (P40 OD010440). H.A.T. is a William Randolph Hearst Investigator. This project was funded in part by a grant from the American Diabetes Association (1-17-IBS-176) to H.A.T. and S.F., a grant from the U.S. Army Research Office (W911NF1210321) to S.F., and an endowment from the William Randolph Hearst Foundation to H.A.T.

## Author Contributions

S.K., Y.S., C.A., S.F. and H.A.T. designed research; S.K., Y.S., C.A., and K.G. performed experiments; S.K., Y.S., and H.A.T. analyzed data; S.K. produced figure models; and S.K. and H.A.T. wrote the manuscript.

## Additional Information (Competing Interests Statement)

None.

