## Supplemental figures for "Bacterial processing of glucose modulates *C. elegans* lifespan and healthspan"

Supplementary Figures and Tables

Samuel F. Kingsley[1α], Yonghak Seo[1α], Calista Allen[2], Krishna S. Ghanta[3], Steven Finkel[2], and Heidi A. Tissenbaum[1,4]*

[1]Department of Molecular, Cell and Cancer Biology, University of Massachusetts Medical School, Worcester, MA 01605

[2]Molecular and Computational Biology Section, Department of Biological Sciences, University of Southern California, Los Angeles, CA, 90089, USA

[33]RNA Therapeutics Institute, University of Massachusetts Medical School, Worcester, MA, 01605, USA

[4]Program in Molecular Medicine, University of Massachusetts Medical School, Worcester, MA, 01605, USA

[α]Co-first author

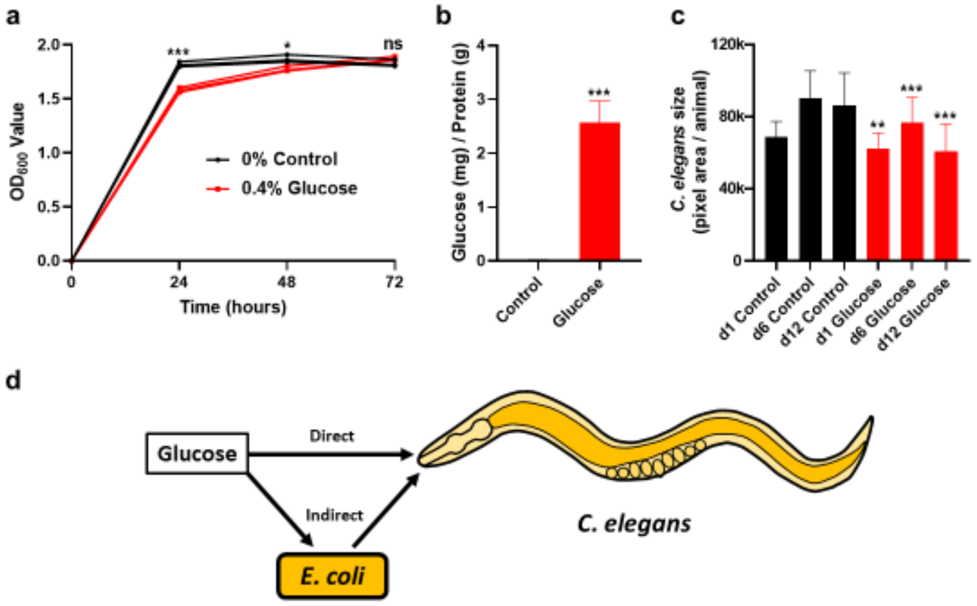


Supplementary Figure S1. Effects of glucose fed bacteria. (a) Optical Density (600nm) of control and 0.4% glucose fed OP50 *E. coli* over 3 days. (b) Glucose assay performed on control and 0.4% glucose fed OP50 *E. coli* after a 3 day culture in LB, results normalized to protein concentration. (c) *C. elegans* body size over time on control and 0.4% glucose fed OP50 *E. coli* measured from photos and quantified by pixel area per animal. (d) Model of direct and indirect bacterial effects of glucose onto its host *C. elegans*.

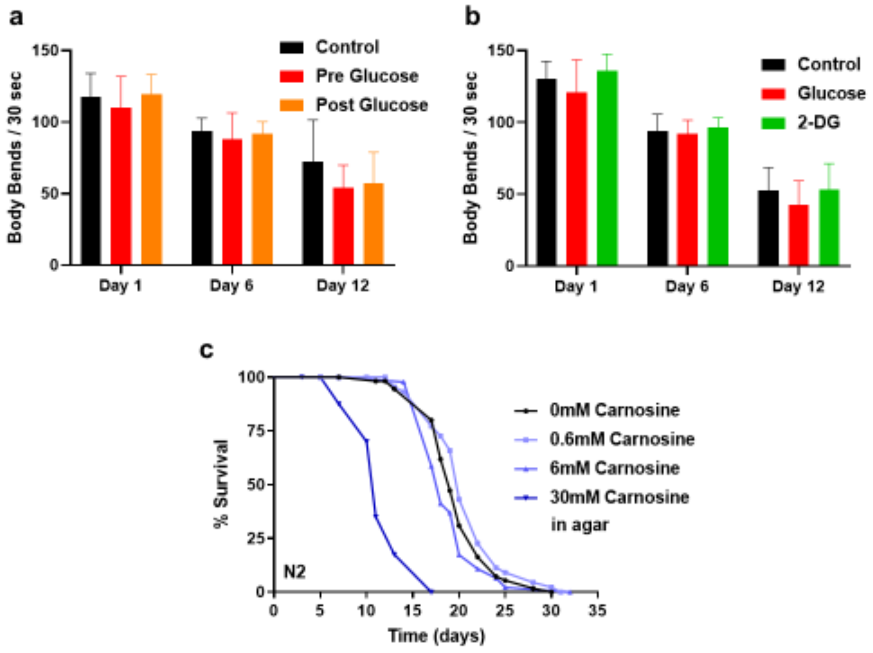


Supplementary Figure 2. Mitigating bacterial metabolism of glucose alters its effects on *C. elegans*. (a) Movement/swimming ability of *C. elegans* treated with control and *E. coli* that was supplemented with 0.4% glucose pre culture or post culture over time. (b) Movement/swimming ability of *C. elegans* treated with 0.4% glucose or 0.4% 2-deoxy-glucose (2-DG) fed OP50 *E. coli* over time. (c) Lifespan assay of wild type *C. elegans* on 0, 0.6, 6, and 30mM Carnosine within the agar seeded with heat killed OP50 *E. coli*.

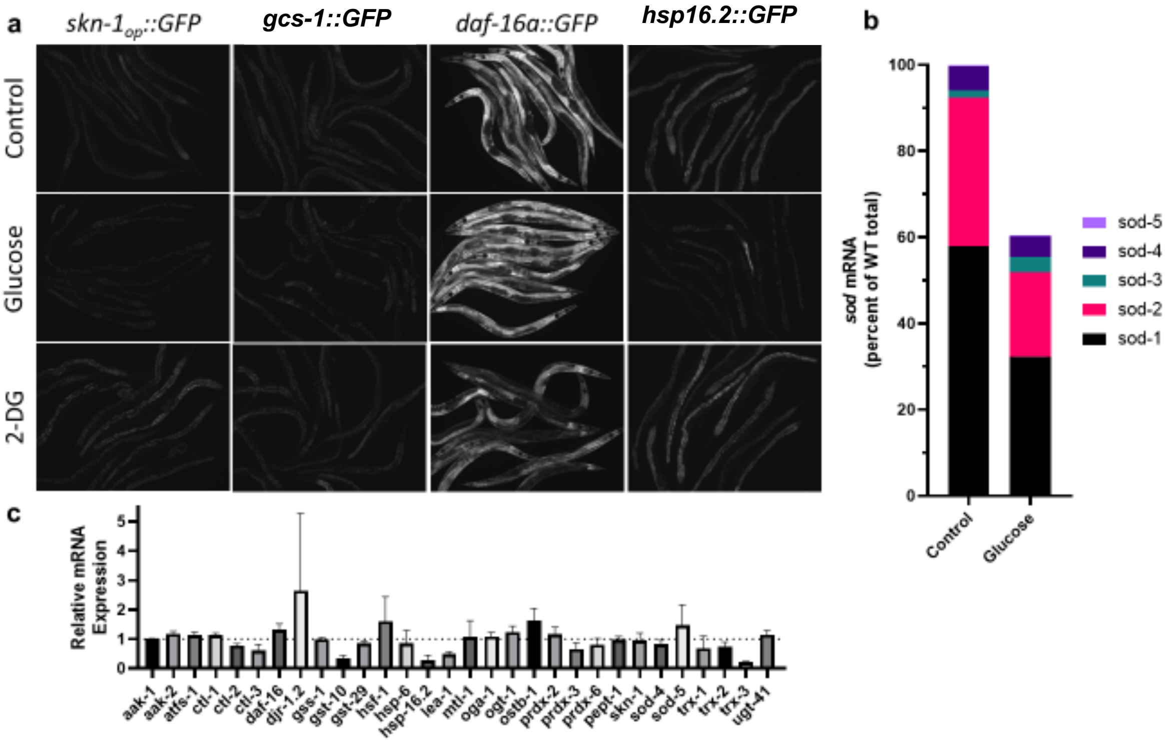


Supplementary Figure 3. *C. elegans* Gene expression changes with bacterial metabolism of glucose. (a) Fluorescent imaging of transgenic strain *C. elegans* [*skn-1*$_{op}$*::gfp*], [*gcs-1::gfp*], [*daf-16::gfp*], or [*hsp-16.2::gfp*], after treatment with control, 0.4% glucose, and 0.4% 2-deoxy-glucose (2-DG) fed OP50 *E. coli* for 6 days. (b) Estimated total *sod* expression within *C. elegans* treated with glucose fed *E. coli* based off the preceding RTqPCR data and published RNA Seq data (Dues et al 2017). (c) RTqPCR experiments performed using wild type *C. elegans* treated with glucose fed *E. coli* for 6 days, relative to expression of *C. elegans* treated with control fed *E. coli*.

| Supplementary Table S1: Lifespan statistics of *C. elegans* on all treatments | | | | | | | |
|---|---|---|---|---|---|---|---|
| **Figure** | ***C. elegans* Genotype** | ***E. coli* Treatment** | **Plate** | **Mean Survival (Days)** | **Std Dev (Days)** | **Animals (N)** | **# Trials** |
| 1D | Wild type (N2) | 0% Control | NGM + Amp | 21.20 | 2.06 | 955 | 12 |
| | | 0.4% Glucose | NGM + Amp | 17.09 | 2.90 | 879 | |
| 2A | Wild type (N2) | 0% Control | NGM + Amp | 21.36 | 1.60 | 361 | 4 |
| | | 0.4% Glucose | NGM + Amp | 17.97 | 2.62 | 345 | |
| | | 0.4% Glucose Post Culture | NGM + Amp | 20.61 | 1.21 | 354 | |
| 2B | Wild type (N2) | 0% Control | NGM + Amp | 21.36 | 0.21 | 275 | 3 |
| | | 0.4% Glucose | NGM + Amp | 16.63 | 2.27 | 165 | |
| | | 0.4% 2-DG | NGM + Amp | 21.55 | 1.18 | 288 | |
| 2C | Wild type (N2) | 0% Control | NGM + Amp | 20.92 | 1.86 | 238 | 3 |
| | | 0.4% Glucose | NGM + Amp | 15.39 | 1.82 | 207 | |
| | | 50mM Carnosine | NGM + Amp | 22.34 | 1.27 | 208 | |
| | | 50mM Carnosine + 0.4% Glucose | NGM + Amp | 19.43 | 1.79 | 244 | |
| 5B | Wild type (N2) | 0% Control | NGM | 19.75 | 2.22 | 273 | 2 |
| | *gst-4(lp11)* | 0% Control | NGM | 16.41 | 0.36 | 146 | |
| | *gst-4(lp10)* | 0% Control | NGM | 17.31 | 0.00 | 90 | 1 |
| 5C | Wild type (N2) | 0% Control | NGM + Amp | 19.72 | 1.40 | 334 | 4 |
| | | 0.4% Glucose | NGM + Amp | 18.39 | 0.51 | 128 | |
| | *gst-4(lp11)* | 0% Control | NGM + Amp | 18.85 | 0.59 | 351 | |
| | | 0.4% Glucose | NGM + Amp | 18.61 | 1.05 | 332 | |

| Supplementary Table S2: RT-qPCR, PCR, and CRISPR primer sequences | | | |
|---|---|---|---|
| **Gene** | **Use** | **Sequence** | **Source** |
| *act-1* | RT-qPCR | CTCTTGCCCCATCAACCATG | Kwon et al 2010 |
| *act-1* | RT-qPCR | CTTGCTTGGAGATCCACATC | Kwon et al 2010 |
| *act-1* | RT-qPCR | GGAGTCATGGTCGGTATGG | GETprime Ensembl release 81 |
| *act-1* | RT-qPCR | CTTGAGGGTAAGGATACCTCTC | GETprime Ensembl release 81 |
| *atfs-1* | RT-qPCR | TTGGAGATAATATGGGCTCCC | GETprime Ensembl release 81 |
| *atfs-1* | RT-qPCR | CTATTCGGGAAGTTCCCGT | GETprime Ensembl release 81 |
| *daf-16* | RT-qPCR | TCCATCATCTTTCCGTCCC | GETprime Ensembl release 81 |
| *daf-16* | RT-qPCR | CTTCCAATAGCTGGAGAAACAC | GETprime Ensembl release 81 |
| *djr-1.1* | RT-qPCR | TTGAGCCATGGAGTCAAGG | GETprime Ensembl release 81 |
| *djr-1.1* | RT-qPCR | AGTACTTGTAGCCTCCTTTCTC | GETprime Ensembl release 81 |
| *djr-1.2* | RT-qPCR | CTGAACCTGTCAAATGTGCC | GETprime Ensembl release 81 |
| *djr-1.2* | RT-qPCR | TGTCGAATGTCTTGTTCTTCAC | GETprime Ensembl release 81 |
| *gcs-1* | RT-qPCR | CCAATCGATTCCTTTGGAGAC | GETprime Ensembl release 81 |
| *gcs-1* | RT-qPCR | CGATGAGACCTCCGTAAGG | GETprime Ensembl release 81 |
| *gss-1* | RT-qPCR | AAGGAAGGATGCACCTGAG | GETprime Ensembl release 81 |
| *gss-1* | RT-qPCR | GCTACTCCACCCATAGCTG | GETprime Ensembl release 81 |
| *gst-10* | RT-qPCR | CTTCACTATTCGAGGATTCGG | GETprime Ensembl release 81 |
| *gst-10* | RT-qPCR | TCGAACCGAATGTCTTCGA | GETprime Ensembl release 81 |
| *gst-29* | RT-qPCR | ATGGAGATGGTACATGGGA | GETprime Ensembl release 81 |
| *gst-29* | RT-qPCR | TAGAACTGGAACTTGGCCA | GETprime Ensembl release 81 |
| *gst-4* | RT-qPCR | GCTGAAGCCAACGACTCCAT | Park, S-K. et al 2009 |
| *gst-4* | RT-qPCR | GACCGAATTGTTCTCCATCGA | Park, S-K. et al 2009 |
| *gst-4* | CRISPR | AAATACAATAGCTTATAGTT | http://crispor.tefor.net |
| *gst-4* | CRISPR | TCGTTGGAGACTCATTGACT | http://crispor.tefor.net |

| | | | |
|---|---|---|---|
| *gst-4* | PCR | TGATGCCAGACGATGACATTAC | IDTdna PrimerQuest tool |
| *gst-4* | PCR | TCTCTGGGAGACGTGATAGG | IDTdna PrimerQuest tool |
| *gst-4* | PCR | CAAATTTCCAGCGACTCCATTT | IDTdna PrimerQuest tool |
| *gst-4* | PCR | ATGATCAGCGTCACTTCCATAG | IDTdna PrimerQuest tool |
| *hsf-1* | RT-qPCR | TCAGACAGTTGAATATGTACGG | GETprime Ensembl release 81 |
| *hsf-1* | RT-qPCR | CCTGATCTGATTCTGTTCGAG | GETprime Ensembl release 81 |
| *hsp-16.2* | RT-qPCR | TGGTGCAGTTGCTTCGAATC | Park, S-K. et al 2009 |
| *hsp-16.2* | RT-qPCR | TTGAACCGCTTCTTTCTTTGG | Park, S-K. et al 2009 |
| *mtl-1* | RT-qPCR | AATCATGGCTTGCAAGTGTG | GETprime Ensembl release 81 |
| *mtl-1* | RT-qPCR | TTCACATTTGTCTCCGCAC | GETprime Ensembl release 81 |
| *oga-1* | RT-qPCR | ACATTATGTGGACAGGACCTC | GETprime Ensembl release 81 |
| *oga-1* | RT-qPCR | GACGCATTACACTTCCCAC | GETprime Ensembl release 81 |
| *ogt-1* | RT-qPCR | TGATCATGACAGGACAAATGAC | GETprime Ensembl release 81 |
| *ogt-1* | RT-qPCR | GATGCATTTGAGACTGTCCG | GETprime Ensembl release 81 |
| *ostb-1* | RT-qPCR | ATGATGTGCAACAGGTGTC | GETprime Ensembl release 81 |
| *ostb-1* | RT-qPCR | CGTAGTATGGATATGCGGAG | GETprime Ensembl release 81 |
| *pept-1* | RT-qPCR | TGCAACACTGGTATTTATGGG | GETprime Ensembl release 81 |
| *pept-1* | RT-qPCR | CGAGATACTTCTCCGAACAC | GETprime Ensembl release 81 |
| *skn-1* | RT-qPCR | GCAACAGCTACTCAATCGT | GETprime Ensembl release 81 |
| *skn-1* | RT-qPCR | TGATGACGAATCAGTAGTGC | GETprime Ensembl release 81 |
| *ugt-41* | RT-qPCR | CTCTTCTAGCTGATTCCCGT | GETprime Ensembl release 81 |
| *ugt-41* | RT-qPCR | TTCCGAGGTAGCTCAACTC | GETprime Ensembl release 81 |

Filename: Supplementary Data 9-4.docx
Directory:
/Users/tissenbh/Library/Containers/com.microsoft.Word/Data/Documents
Template: /Users/tissenbh/Library/Group Containers/UBF8T346G9.Office/Library/Normal.dotm
Title:
Subject:
Author: Kingsley, Samuel
Keywords:
Comments:
Creation Date: 9/4/20 4:19:00 PM
Change Number: 11
Last Saved On: 9/13/20 7:48:00 PM
Last Saved By: Tissenbaum, Heidi A.
Total Editing Time: 39 Minutes
Last Printed On: 9/13/20 7:50:00 PM
As of Last Complete Printing
Number of Pages: 7
Number of Words: 914
Number of Characters: 5,737 (approx.)
