## Supplementary figures and images for "Bacterial processing of glucose modulates *C. elegans* lifespan and healthspan"

### Supplemental data

Supplemental
Figure 1

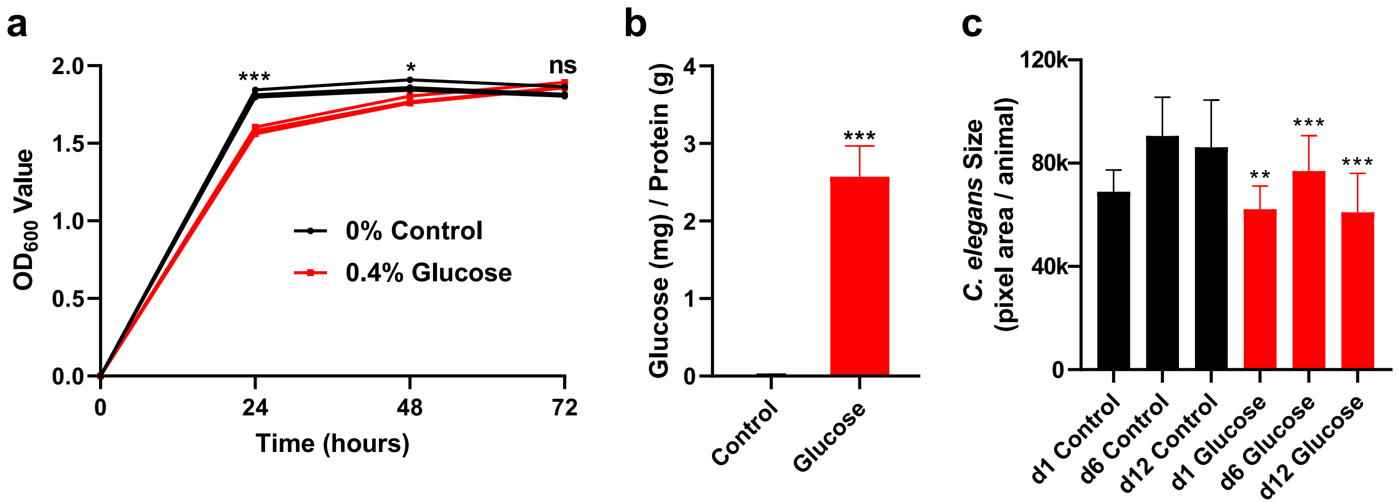

Supplemental
Figure 2

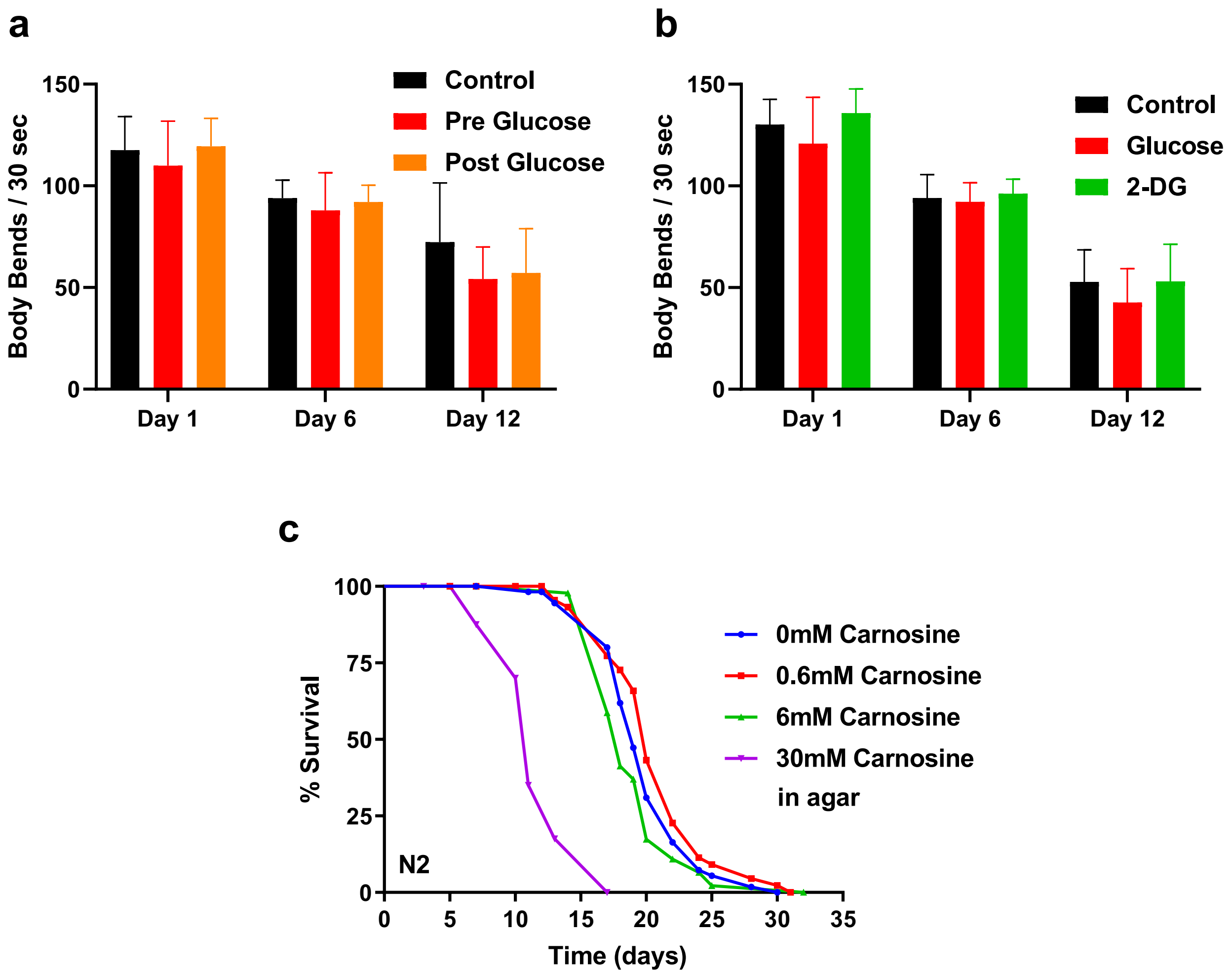

Supplemental Figure 3

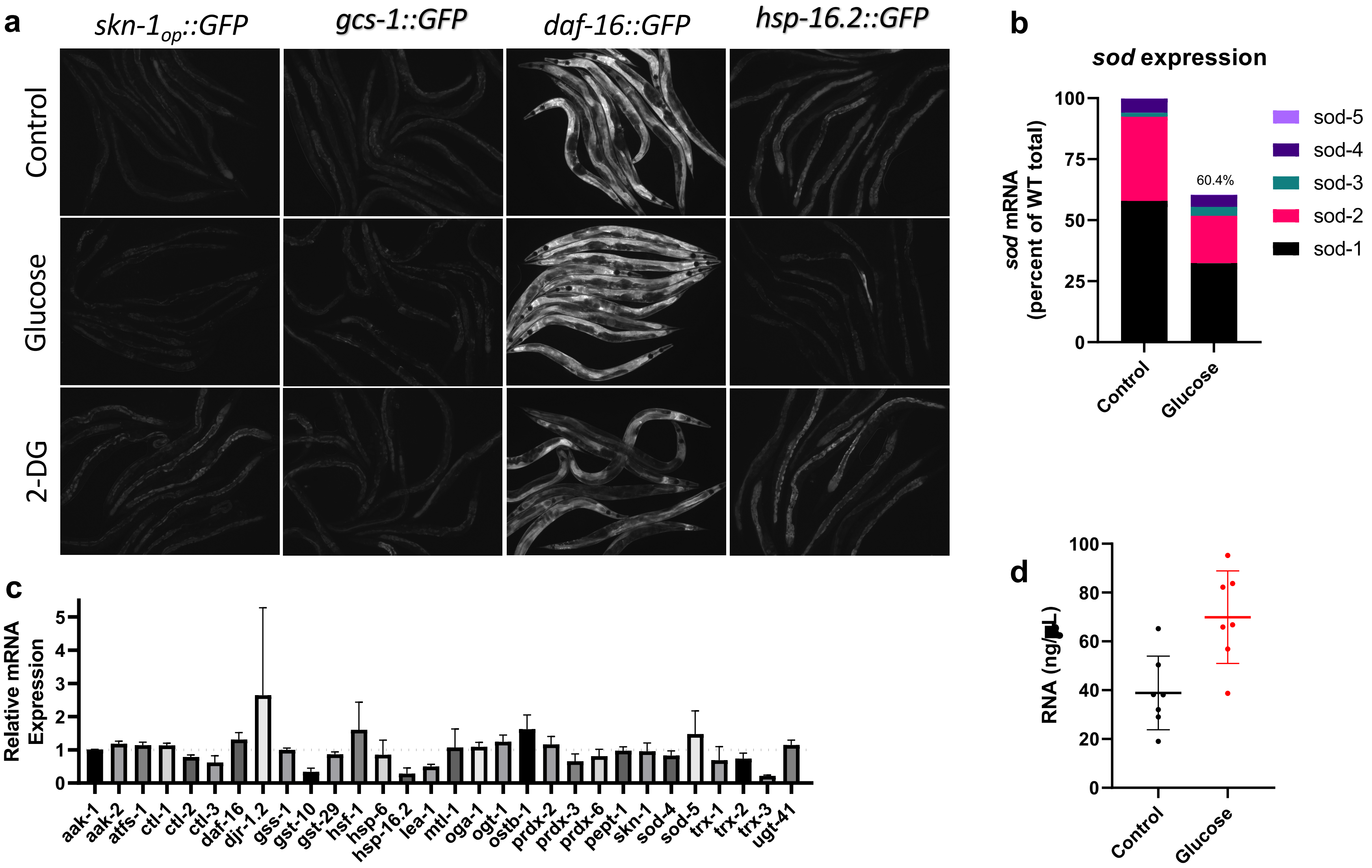
